# Presence of Hepatitis B virus infection in colon cancer

**DOI:** 10.1101/2025.03.30.646256

**Authors:** Lishi Li, Wenjing Gong, Yunyun An, Hua Tong, Mengqi Yang, Wanqiu Wang, Li Ma, Shiliang Tu, Yang Liu, Yunfeng Duan, Kun Sun

## Abstract

Hepatitis B virus (HBV) infection affects ∼300 million people worldwide, which population is at an elevated risk of various cancers, such as gastrointestinal cancer, while the matter behind is largely unclear. In this study, we analyzed 10 colonic specimens from 6 colon cancer patients who were also chronic HBV carriers through Fluorescence-activated Cell Sorting and Immunofluorescence assay to inspect the expression of HBsAg (HBV surface antigen), NTCP (current known cellular receptor of HBV) and CD45 (common marker for immune cells) proteins in single-cell resolution. We found that presence of HBsAg in colonic cells was observed in specimens from all patients, and the proportions of HBV-infected cells were higher in tumors than normal tissues. Strikingly, HBsAg in both immune cells and NTCP-negative cells were also observed in all the patients. *In vitro* experiments further demonstrated that colonic cells expressing NTCP were indeed highly suspectable to HBV infection. Hence, HBV could infect colonic cells (including immune ones) and may play roles during tumorigenesis in colon cancer.

## Introduction

Hepatitis B virus (HBV) is one of the most common infectious viruses in human. WHO (World Health Organization) estimates that there are as many as 300 million people suffering from chronic HBV infection worldwide; WHO Western Pacific Region (including China) and African Region possess the highest rate of HBV infection ^1^. Clinically, infection of HBV is routinely diagnosed if the tests for HBV-related antigens (e.g., Hepatitis B virus Surface antigen (HBsAg)) and HBV-DNA show positive results in the patients’ peripheral blood. HBV is considered as a hepatotropic virus and mainly infects the liver tissue. At present, NTCP (Sodium taurocholate cotransporting polypeptide; the official gene symbol is *SLC10A1*) is the only functionally validated entry receptor of HBV ^2^, and many studies (e.g., the Genotype-Tissue Expression (GTEx) project ^3^) have shown that NTCP is highly expressed in hepatocytes while barely expressed in other human cells. Nonetheless, recent studies have revealed that HBV carriers are at a much higher risk of various cancer types besides liver cancer ^4,5^. For instance, it is reported that the morbidity of gastrointestinal cancer in HBV carriers is more than 5-fold higher than non-HBV carriers in China ^5^. However, the underline mechanisms linking HBV infection and gastrointestinal neoplasms are largely elusive.

Gastrointestinal cancer is a general designation for cancers affecting the gastrointestinal tract; cancers in the gut (e.g., colorectal carcinoma) are the most common types of gastrointestinal cancer and are also among the top-ranked cancers with high morbidity and mortality ^6^. The gut is known as a reservoir for miscellaneous types of microbiomes, including bacteria and viruses. The compositions, functions and relationships with various diseases of the gut bacteriome have been relatively well profiled ^7^; as a contrast, although studies have suggested that viruses in the gut may involve in carcinogenesis of colorectal carcinoma ^8^, however, the gut virome remains largely uncharacterized. In particular, it is reported that in a subset of chronic HBV carriers, the virus could be detected in the colonic mucosa ^9^; however, considering the elevated morbidity rate of gastrointestinal cancer in HBV carriers, further explorations on HBV and colon cancer, such as whether HBV could infect colonic cells, had not been convincingly performed. To this end, in this study, we utilized various cell analysis techniques to investigate the co-existence of HBV and its receptor, NTCP, in colonic cells.

## Results

### Overview of the study

Table 1 summarized the patients, specimens, experiments, as well as major results of this study. Briefly, 6 colon cancer patients were recruited (referred to as patients P1P6 hereafter; detailed information were provided in Suppl. Table S1) in this study; both the tumor samples and adjacent normal tissues were collected from P1, P2, P4, and P6, while only tumor samples were available for P3 and P5. Fluorescence-activated Cell Sorting (FACS) was performed on all specimens to investigate the presence and proportion of HBsAg-positive cells. We further utilized multiplexed immunofluorescence assay to validate the co-localization of HBsAg, NTCP and CD45 (common surface marker for immune cells; the official gene symbol is *PTPRC*) in single-cell resolution on specimens from patient P2, P3, P5, P6, as specimens from P1 and P4 were too limited such that we did not have any samples left after FACS.

**Table 1.** Summary of specimens, experiments, and major results.

| Patient |  | P1 |  | P2 |  | P3 |  | P4 |  | P5 | P6 |  |
| --- | --- | --- | --- | --- | --- | --- | --- | --- | --- | --- | --- | --- |
| Specimen |  | T | N | T | N | T |  | T | N | T | T | N |
| FACS | CD45-NTCP+HBV+ | + | - | + | + | + |  | + | + | - | - | - |
|  | CD45-NTCP-HBV+ | + | + | + | - | + |  | + | + | + | - | - |
|  | CD45+NTCP+HBV+ | + | + | + | + | + |  | + | + | + | + | + |
|  | CD45+NTCP-HBV+ | + | + | + | + | - |  | + | + | + | + | + |
| IF | CD45-NTCP+HBV+ |  |  | + | + | + |  |  |  | - | - | - |
|  | CD45-NTCP-HBV+ | na |  | + | + | + |  | na |  | - | - | - |
|  | CD45+NTCP+HBV+ |  |  | + | + | + |  |  |  | + | + | + |
|  | CD45+NTCP-HBV+ |  |  | + | + | + |  |  |  | + | - | + |

Note: '+' and '-' stands for detectable and undetectable, respectively; 'na' means the experiment was not performed.

### Fluorescence-activated Cell Sorting (FACS) results

We first investigated the presence and proportions of HBsAg-positive cells in all specimens. As shown in Table 1, Fig. 1, and Suppl. Fig. S1, HBsAg-positive cells were detected in all patients with variable proportions. For instance, as shown in Fig. 1, 5.2% and 78.9% CD45-negative cells (i.e., non-immune cells, which category contains the malignant/cancerous cells in tumors) were positive for both NTCP and HBsAg in the tumors of P1 and P2, respectively; in contrast, NTCP and HBsAg double-positive, CD45-negative cells were not detectable in the adjacent normal tissue of P1, while detectable in P2 with a much lower fraction (10.1%) compared to the tumor. Surprisingly, in the CD45-negative cells from all specimens, a remarkable population of HBsAg-positive cells were negative for NTCP (i.e., these cells did not express the known HBV receptor). In addition, in all specimens, an extra-high proportion of CD45-positive cells (i.e., infiltrating immune cells) was also positive for HBsAg; in these CD45- and HBsAg-double-positive cells, a remarkable proportion were negative for NTCP, which observation was similar to that of CD45-negative cells. These results thus suggested that colonic cells, including the immune ones, may get infected by HBV, even without the expression of NTCP. In the meantime, in P1, P2, P4, which patients had paired tumor and adjacent normal samples and detectable malignant cells with potential HBV infection (i.e., CD45-negative, NTCP and HBsAg double-positive cells), the proportions of such cells were all higher in tumors compared to adjacent normal tissues. Moreover, single-cell RNA-seq data on colorectal cancer samples validated the expression of NTCP in both immune and non-immune cells (Suppl. Fig. S2).

**Fig. 1.**
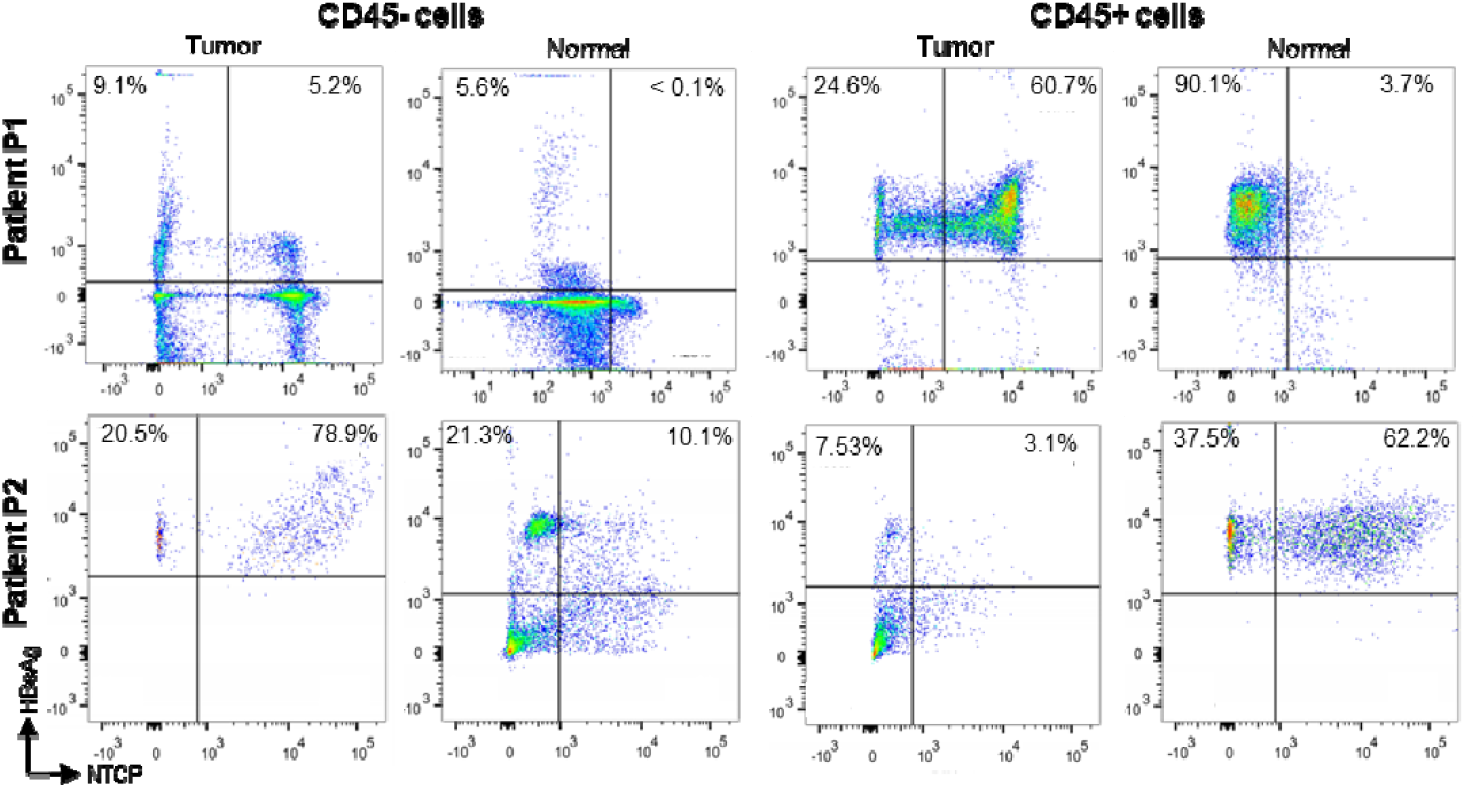
Flow cytometry results on colonic specimens from patient P1 and P2. The CD45-negative cells contain the malignant cells in tumor; CD45-positive cells are infiltrating immune cells.

**Fig. 2.**
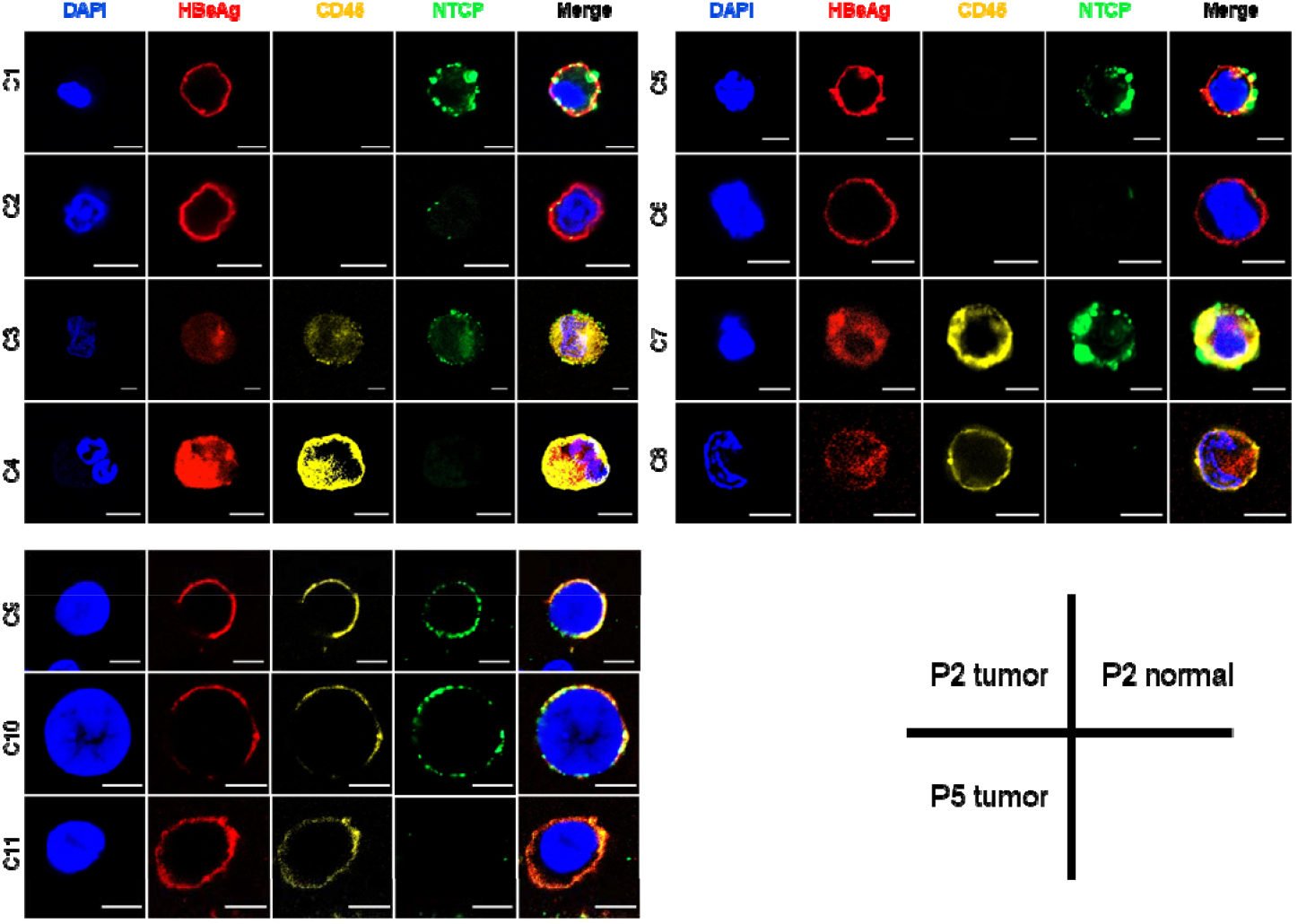
Representative cells from single-cell immunofluorescence experiment on specimens from patient P2 and P3. C1-C4 were cells from the tumor sample of patient P2; C5-C8 were cells from the adjacent normal tissue of patient P2; C9-C13 were cells from the tumor sample of patient P5. Scale bars indicated 5um.

### Immunofluorescence staining

To validate the results by flow cytometry, we utilized multiplex immunofluorescence staining and imaging analysis to explore the spatial co-localization of HBsAg, NTCP, and CD45 in dissociated single cells from patients P2-P6. Representative imaging results for HBsAg-positive cells were shown in Figure 2 and Suppl. Fig. S3: cells C1-C4, and C5-C8 were originated from the tumor sample and adjacent normal tissue of P2, respectively; cells C9-C12 were from the tumor sample of P3. C1, C2, C5, C6, C9, and C10 were CD45-negative while the rest were CD45-positive. Notably, C2, C4, C6, C8, C10, and C12 showed strong HBsAg signal, while either very weak or no NTCP signal. The data confirmed the presence and various types of HBsAg-positive cells in colon. In addition, the Human Protein Atlas database ^10^ provided NTCP antibody staining results for 2 colon samples from different donors, and NTCP expression was readily detectable in glandular cells in both samples (Suppl. Fig. S4), which was consistent with our results.

### In vitro infection

We found that expression of NTCP could be detected in 4 commonly used colonic cell lines both in mRNA and protein level (Fig. 3a-b), which results were consistent with the transcriptome data in Cancer Cell Line Encyclopedia (CCLE) database ^11^. Using FACS, we further validated the expression of NTCP protein on the cell membrane of Caco2, which showed the highest expression of NTCP in the 4 cell lines investigated (Suppl. Fig. S5), and collected cells that expressed NTCP (named as “Caco2-NTCP+” hereafter; Suppl. Fig. S6). We then investigated the susceptibility of Caco2 cells to HBV entry. To do this, we engineered a pseudovirus using a HIV backbone and HBV-L protein, which is responsible for binding to NTCP and facilitating HBV entry, was expressed and displayed on the surface of the virual particles (Suppl. Fig. S7a-b). Additionally, we introduced the firefly luciferase gene into the HBV-L pseudovirus genome (HBV-L luciferase) to quantify the HBV entry capability into Caco2 cells. The HBV-L luciferase pseudoviruses were successfully produced and verified by HIV p24 protein quantification, and the HBV L protein was abundantly expressed on these HBV-L pseudoviruses (Suppl. Fig. S7c-d).

**Fig. 3.**
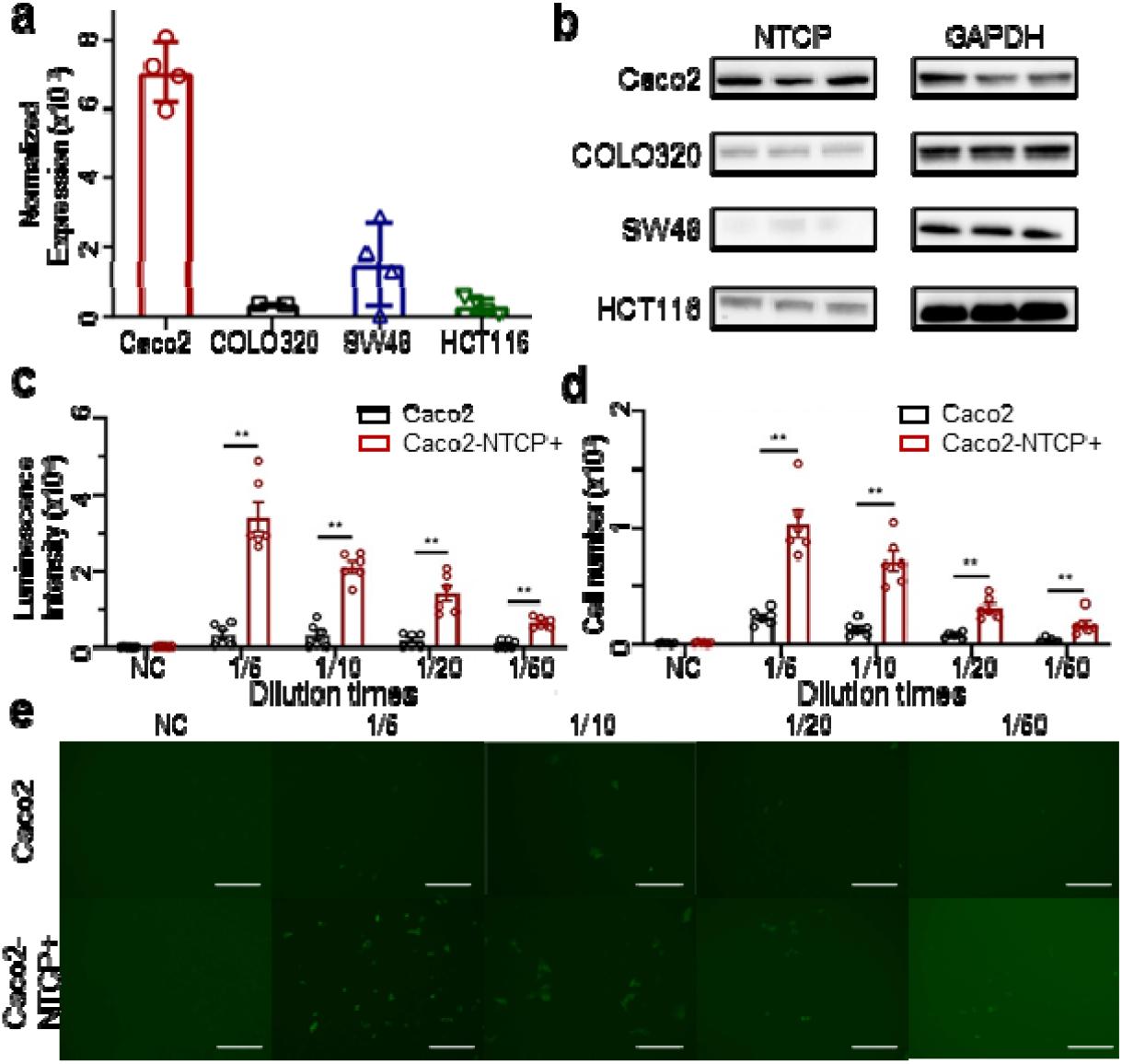
Expression of NTCP in colonic cells and potential for HBV infection. (a) mRN expression (normalized by GAPDH) of NTCP in 4 colon cancer cell lines using qPCR. (b) Western Blot results of NTCP expression in 4 colon cancer cell lines. (c-e) Infection of HBV-L pseudovirus into wildtype Caco2 and Caco2-NTCP+ cells. (c) Luciferase and (d) mNeonGreen fluorescence signals generated by the entered HBV-L genome. (e) Representative images of mNG-positive cells from two independent repeats. Scale bar: 300 μm. Dots on the graph represent individual biological replicates pooled from two independent experiments. The values in the graph represent the mean ± standard error of mean. Statistical analysis were performed using two-tailed Mann-Whitney tests, **: p < 0.01, ***: p < 0.001. NC: negative control.

We exposed wildtype Caco2 and Caco2-NTCP+ cells to serially diluted HBV-L luciferase pseudoviruses for 48 hours. Then, the cells were lysed, and firefly luciferase signal was detected using the Bright-light luciferase assay system. The luciferase intensities were 6.9 to 10.5 times higher in Caco2-NTCP+ cells across various initial infection pseudovirus concentrations when compared to Caco2 wildtype cells (Fig. 3c), suggesting that Caco2-NTCP+ cells with elevated NTCP protein expression exhibit an enhanced susceptibility to HBV pseudovirus entry. To visually observe HBV-L pseudovirus infection in Caco2 cells, we replaced the luciferase reporter gene with the mNeonGreen fluorescence gene and produced the new HBV-L mNeonGreen (HBV-L mNG) pseudovirus (Fig. 3d-e). The HBV-L mNG pseudovirus also attains a high yield of HBV L protein expression (Supplementary Fig. S7d). We inoculated HBV-L mNG pseudovirus onto both Caco2 and Caco2-NTCP+ cells. The number of infected cells displaying green fluorescence was calculated 4 days post-infection. As expected, a greater number of infected cells were observed in Caco2-NTCP+ cells in a dose-dependent manner (Fig. 3d). These results thus demonstrated that Caco2 cells possessed high potential to be infected by HBV.

## Discussion

In this study, we report the existence of HBsAg-positive cells in colonic samples. In particular, we observed high proportions of HBsAg and NTCP double-positive cells in all the 6 patients investigated, which strongly suggests the presence of HBV infection in colon cancer. NTCP is the only validated entry receptor of HBV at present; besides its biased expression in liver, data from previous studies do support the expression of NTCP in colon tissue, both in RNA and protein levels (Suppl. Fig. S2, S4). In addition, NTCP is also expressed in various colonic cancer lines, and these NTCP-expressing colonic cells are highly susceptible to HBV infection (Fig. 3). Hence, these relevant and consistent data provide ponderable supports for our results and conclusions.

Interestingly, we observed that a high proportion of tissue infiltrating immune cells (i.e., CD45-positive cells) are also positive for NTCP and/or HBsAg. One possible explanation is that these immune cells get HBsAg though phagocytosis of dead HBV-infected malignant cells (i.e., they are not actually infected by HBV) ^12^. However, we could not rule out the possibility that the immune cells are actually get infected. In fact, a recent study on COVID-19 suggests that SARS-CoV-2 might be able to infect immune cells even though the immune cells do not express the entry receptor (i.e., ACE2) of SARS-CoV-2 ^13^, which situation is similar to our observation that HBsAg-positive while NTCP-negative cells do exist. Hence, although our data showed that NTCP was expressed in colonic cells and such NTCP-expressing cells were suspectable to HBV infection, it could also be possible that HBV could infect colonic cells (including infiltrating immune cells) without NTCP ^14^, while further comprehensive molecular studies are necessary to answer this essential question.

In addition, our data show that in all 4 patients with paired tumor and adjacent normal samples, the proportion of HBV-infected cells are higher in the tumor than normal samples (Table 1), suggesting the possibility that HBV is involved in tumorigenesis of colon cancer. This hypothesis could (partially) explains the observation that HBV carriers are at a much higher risk of gastrointestinal cancers ^5^. Biologically, there are also lots of directions worthwhile for further investigations, especially the molecular mechanism of HBV in tumorigenesis. In clinical side, however, there are lots of questions to be answered. For instance, how many colon cancer patients suffer from HBV-infection in colonic cells? How does HBV-infection affect the therapeutics and prognosis of these patients? We think that these questions are very important and may affect clinical practices in colon cancer treatment, therefore worthwhile for large-scale investigations.

## Methods

### Ethics approval and sample collection

This study had been approved by the Ethics Committee of Zhejiang Provincial People’s Hospital and Ethics Committee of Shenzhen Bay Laboratory. A total of 6 patients were recruited from Zhejiang Provincial People’s Hospital and written-informed consents had been obtained from all participants. Basic information of these patients could be found in Suppl. Table S1. All the 6 patients were chronic HBV carriers as determined by routine blood tests. After obtaining the colon tumors and adjacent normal tissues (if any) during surgical resections, the specimens were immediately washed using physiological saline, then stored in MACS Tissue Storage Solution (Miltenyi Biotec, #130-100-008) and analyzed within 48 hours.

### Tissue dissociation and single cell suspension preparation

Colon tumors and adjacent normal tissues were cut into small pieces and digested with Tumor Dissocation Kit, human (Miltenyi Biotec, #130-095-929) in gentleMACS C Tubes (Miltenyi, #130-093-237) following the manufacturer’s protocol. The tubes were then placed on a gentleMACS Octo Dissociator with Heaters (Miltenyi, #130-096-427), and the built-in program ‘37C_h_TDK_1’ was run for 1 hour to digest the samples. Digested samples were filtered using 100 μm and 40 μm cell strainers and diluted with RPMI 1640 medium (Gibco, #11875176). Cells were centrifuged at 400g at 4°C for 10 min. Red blood cell lysis was performed with Red Cell Lysis Buffer (TIANGEN, #4992957). Remaining cells were resuspended with 0.04% BSA in PBS (phosphate buffer saline).

### Cell staining and flow cytometry analysis

Cells were then stained in cell staining buffer (Biolegend, #420201) with a combination of fluorescent antibodies: Alexa Fluor 700 anti-human CD45 (1:20; Biolegend, #304024), NTCP Polyclonal Antibody (1:100; Thermo Fisher, #PA5-25614). Intracellular staining of HBsAg (Hepatitis B virus Surface antigen) was performed using Hepatitis B virus Surface Monoclonal Antibody (S26; 1:100; Invitrogen, #MA1-7603) and the Cytofix/Cytoperm kit (BD Biosciences, #554714) according to the manufacturer’s protocol. The stained cells were analyzed using a BD FACSAria III flow cytometer, and the resulting data was analyzed using FlowJo (v10.4.1, Tree Star). We first separated the cells into CD45-negative and CD45-positive; then for each category (e.g., NTCP+HBV+), we consider it “detectable” if the cells in this category accounted for more than 1% of all the cells.

### Immunofluorescence assay

Immunofluorescence was performed using conventional protocols with minor modifications ^15^. For single-cell immunofluorescence, single cell suspension was prepared using Tumor Dissociation Kit, human (Miltenyi Biotec, #130095929). For tissues, fresh tissue was embedded with Optimal Cutting Temperature Compound (SAKURA, #4583). Tissue blocks were sliced into 15 μm and rinsed with PBS. Single cells and histology slides were fixed with 4% paraformaldehyde (Biosharp, #BL539A) and followed by PBS rinsing. 0.1% Triton X-100 were used to permeabilize cells and unspecific binding were blocked with 5% BSA in PBS. Single cells and slides were incubated with Alexa Fluor 700 anti-human CD45 (Biolegend, #304024), NTCP Polyclonal primary Antibody (Thermo Fisher, #PA5-25614) and Hepatitis B virus Surface Monoclonal Antibody (S26; Invitrogen, #MA1-7603) and followed by washing with 0.1% Tween-20 in PBS. Then single cells and slides were incubated with secondary antibodies as followings: AF488-conjugated anti-mouse IgG (1:2000; Cell Signaling Technology, #4408) and APC-conjugated anti-mouse IgG (1:2000; Cell Signaling Technology, #4414). Single cells and slides were mounted with 5 ul ProLong Gold Antifade Reagents with DAPI (Invitrogen, #P36930) and sealed with nail polish. Stained slides were digitalized on an LSM 900 confocal microscope (ZEISS, Germany), and the imaging data was processed using the ZEISS ZEN software (v3.2, blue edition) with Airyscan detector.

### Cell culture

Caco2, HCT116, SW48 and COLO320 cells were purchased from BNCC. All cells were cultured in Dulbecco’s modified eagle medium (DMEM; Thermo Fisher, #C11995500BT) supplemented with 10% fetal bovine serum (FBS) and 100 U/mL penicillin/streptomycin (Corning, #30-002-CI) at 37 °C in the humidified incubator with 5% CO_2_. All experiments were performed with Caco2 between passages 2–10. Mycoplasma contaminations were tested every week using the GMyc-PCR Mycoplasma Test Kit GMyc-PCR (Yeasen, #40601ES10).

### Western blotting

Western blotting was performed using standard protocols. In brief, cells were harvested and lysed in the RIPA lysis buffer (Beyotime, #P0013) containing protease inhibitor (Roche, #5892953001) on ice for 30 min. Samples were centrifuged and the supernatant was collected, followed by quantification with Detergent Compatible Bradford Protein Assay Kit (Beyotime, #P0006C). Proper amounts of proteins were separated in 4-12% gradient SDS-PAGE gels (GenScript, #M00653) and transferred to PVDF membranes (Millipore, #IPVH00010). Membranes were blocked in 5% defatted milk and incubated with anti-NTCP primary antibody (1:1000; Thermo Fisher, # PA580001) or HRP Conjugate anti-GAPDH antibody (1:3000; Cell Signaling Technology, #8884) overnight at 4°C. Membranes were washed with 0.1% Tween-20 in TBS. The membrane incubated with anti-NTCP primary antibody was then incubated with HRP-linked Anti-rabbit IgG (1:5000; Cell Signaling Technology, #7074S) and followed by washing with 0.1% Tween-20 in TBS. Proteins were visualized with Enhanced ECL Chemiluminescent Substrate Kit (Yeasen, #36222es76) and Bio-Rad ChemiDoc Imaging Systems (Bio-Rad, USA).

### RNA Isolation, Reverse Transcription, and quantitative RT-PCR

RNA was isolated with TRIzol Reagent (Thermo Fisher, #15596018) and Direct-zol RNA kit (Zymo, #R2052) following the manufacturer’s instructions. 1 μg RNA was reverse transcribed using the HiScript III All-in-one RT SuperMix Perfect for qPCR (Vazyme, #R333) following the manufacturer’s instructions. Relative gene expression levels were analyzed by quantitative RT-PCR (qRT-PCR) with the Taq Pro Universal SYBR qPCR Master Mix (Vazyme, #Q712) on a QuantStudio 3 Real-Time PCR System (Thermo Fisher, USA). The primers involved were as follows:

*SLC10A1*:

forward 5’- GGACATGAACCTCAGCATTGTG -3’,

reverse 5’- ATCATAGATCCCCCTGGAGTAGAT -3’.

*GAPDH*:

forward 5’- AGGTCGGTGTGAACGGATTTG -3’,

reverse 5’- GGGGTCGTTGATGGCAACA -3’.

### Sorting NTCP+ cells in Caco2 cell line

Caco2 were harvested when the confluence reached to 70%-80% and stained in cell staining buffer (Biolegend, #420201) with NTCP Polyclonal Antibody (Thermo Fisher, #PA5-25614) and Donkey anti-Rabbit IgG (H+L) Highly Cross-Adsorbed Secondary Antibody, Alexa Fluor Plus 488 (Thermo Fisher, #A32790). The stained cells were analyzed using a Beckman CytoFLEX SRT Benchtop Cell Sorter, and the resulting data was analyzed using FlowJo (v10.4.1, Tree Star). Caco2-NTCP+ cells were cultured under the same condition as Caco2 wildtype cells.

#### Packaging of HBV-L pseudovirus

HBV-L Luciferase Pseudovirusor HBV-L mNeonGreen (mNG) Pseudovirus were generated by co-transfecting the HBV L-encoding plasmid and the human immunodeficiency virus (HIV) backbone expressing firefly luciferase (pNL4-3.luc.RE) or mNeonGreen (pNL4-3.mNeonGreen, modified from pNL4-3.luc.RE) into 293T cells (ATCC, #CRL3216). The transfection was carried out using Lipo8000™ Transfection Reagent (Beyotime, #C0533) following the manufacturer’s instructions. Pseudotyped virus stocks were collected 48 hours after transfection, filtered, and stored at -80°C. Viral titers were determined using a p24 ELISA kit (SinoBiological, #KIT11695) according to the manufacturer’s instructions.

### Pseudovirus infection and luciferase detection

For pseudovirus infection, 1.0×10^4^ Caco2 cells were seeded in 100 μL of DMEM containing 2% FBS and 100 U/mL Penicillin-Streptomycin (P/S) in each well of a flat-bottom 96-well plate. After 16 hours culture, serially diluted pseudoviruses in 125 μL of DMEM containing 2% FBS (OmnimAbs, USA) and 100 U/mL P/S was added to each well. The cells were then incubated in a 5% CO2 environment at 37°C for 48 hours. HBV-L Luciferase pseudovirus infection was evaluated by quantifying luciferase activity with the Bright-Lite Luciferase Assay System (Vazyme, #DD1204) using a multifunctional microplate detector (TECAN, Switzerland).

### Quantification and imaging of HBV-L mNG infected cells

The procedures for HBV-L mNeonGreen (HBV-L mNG) pseudovirusinfection were similar to those for HBV-L Luciferase pseudovirus. The cell status was monitored daily, and infected cells displaying green fluorescence were imaged using a Cell Imaging System (Invitrogen EVOS M5000, USA) 4 days post-infection. The number of infected cells was quantified using a BioTek Cytation 7 Cell Imaging Multimode Reader (Agilent, USA). Twenty photos per well were taken using the GFP channel under 4× phase and combined into a single image for each well. Cell counting was performed using the cellular analysis program of Gen5 3.04 software according to the manufacturer’s instructions.

## Acknowledgement

This study was supported by National Key R&D Program of China (2022YFA0912700 to K.S.), National Natural Science Foundation of China (NSFC; 32170068 to Y.D.), Guangdong Basic and Applied Basic Research Foundation (to KS), and Shenzhen Bay Laboratory (to Y.L. and K.S.). We would like to thank Ms. Qi Wang, Shenzhen Bay Laboratory, for her technical assistances; SZBL Biochemistry Core and Imaging Core for providing flow cytometry and imaging supports, respectively.

## Author contributions

W.G., Y.D., K.S. conceived and designed research; Y.L., K.S. supervised research; L.L., Y.A., HT, M.Y., W.W., L.M. performed research; L.L., Y.A. HT, Y.L., Y.D., K.S. analyzed data; S.T., W.G. recruited subjects and analyzed clinical data; L.L., Y.A., Y.L., Y.D., K.S. wrote the paper.

