## Supplementary material for "Presence of Hepatitis B virus infection in colon cancer": Supp. Tables and Figures

Li *et al.*

This supplementary information contains Table S1 and Figures S1-S7.

**Table S1. Basic information of the patients.**

| Patient ID | Age | Gender | Tumor | Adjacent Normal |
|---|---|---|---|---|
| P1 | 59 | M | Yes | Yes |
| P2 | 48 | M | Yes | Yes |
| P3 | 56 | F | Yes | No |
| P4 | 56 | F | Yes | Yes |
| P5 | 55 | F | Yes | No |
| P6 | 58 | F | Yes | Yes |

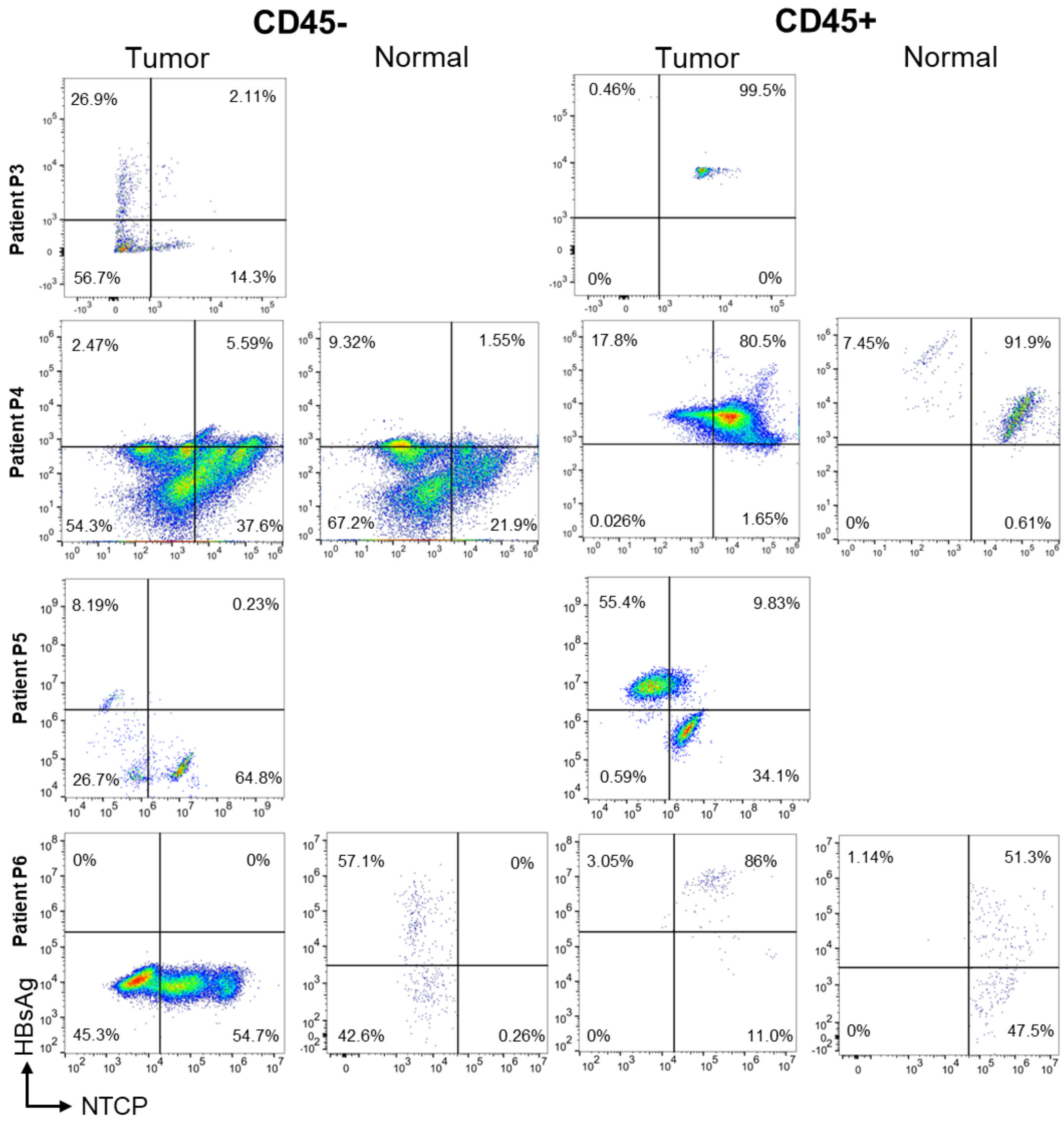


**Fig. S1. Flow cytometry results on colonic specimens from patients P3 to P6**. The CD45-negative cells contain the malignant cells in tumor; CD45-positive cells are infiltrating immune cells.

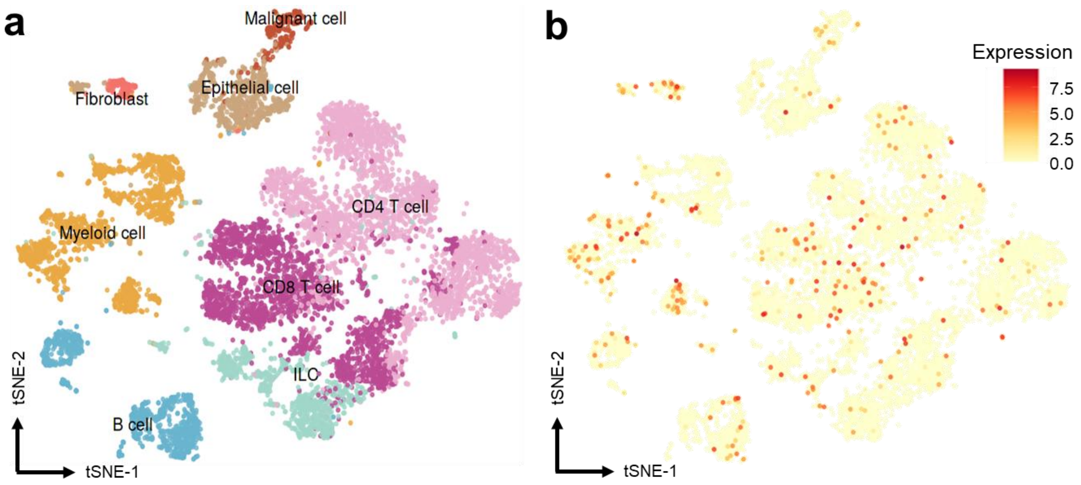


**Fig. S2. SLC10A1 expression in scRNA-seq of colon samples**. (a) Cell clusters and annotations. (b) Expression levels for SLC10A1.

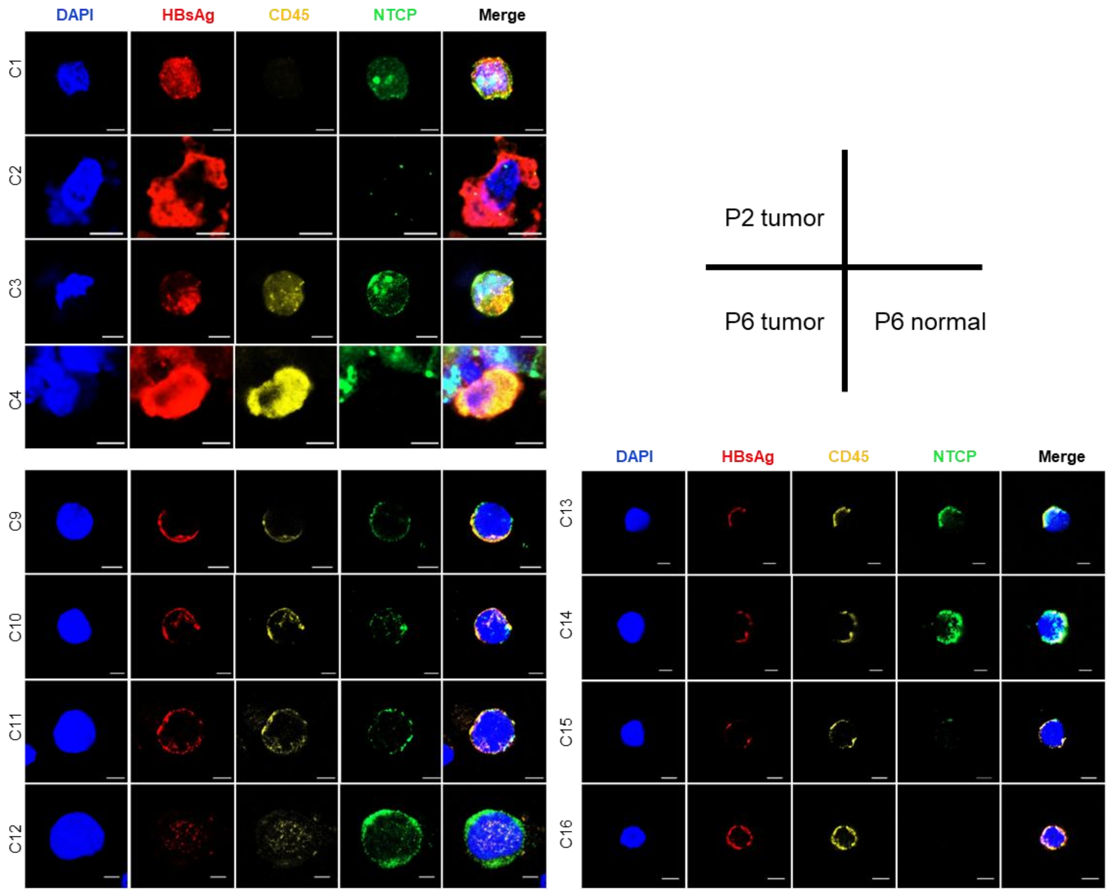


**Fig. S3. Representative cells from single-cell and tissue immunofluorescence experiment on specimens from patients P5 and P6.** C1-C4 were cells from the tumor sample of patient P2; C5-C8 were cells from the tumor of patient P6; C9-C12 were cells from the adjacent normal tissue sample of patient P6. Scale bars indicated 5um.

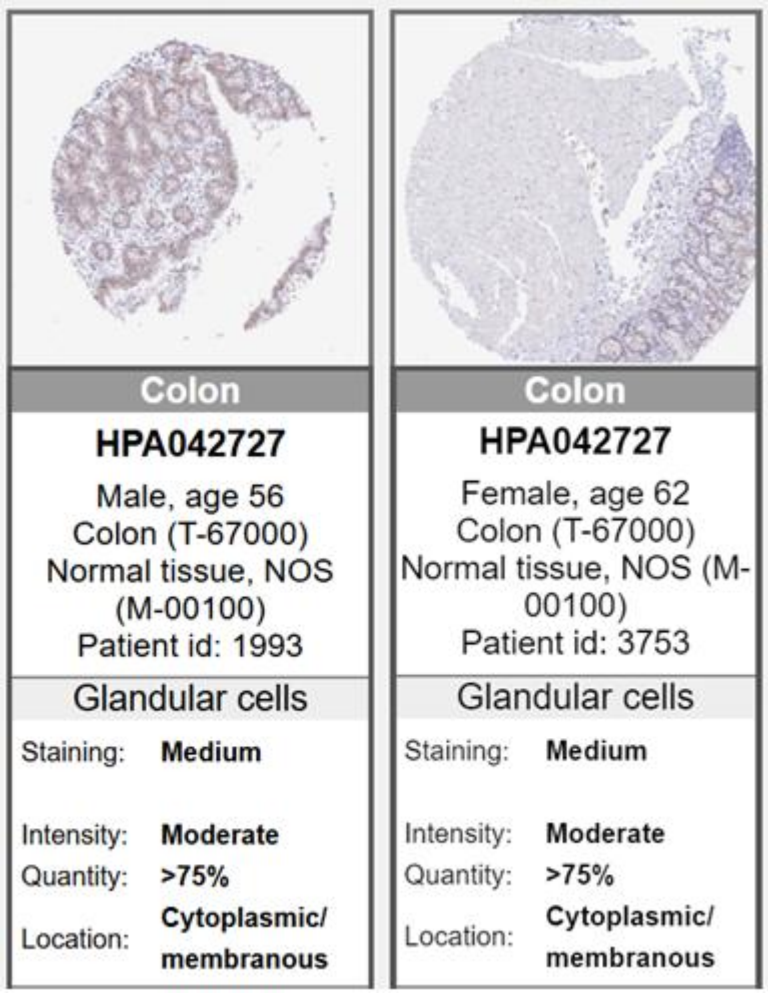


**Fig. S4. Immunohistology chemical of NTCP in colon.** The data is obtained from the Human Protein Atlas (https://www.proteinatlas.org/ENSG00000100652-SLC10A1/tissue/colon), fetched on Oct 1, 2023.

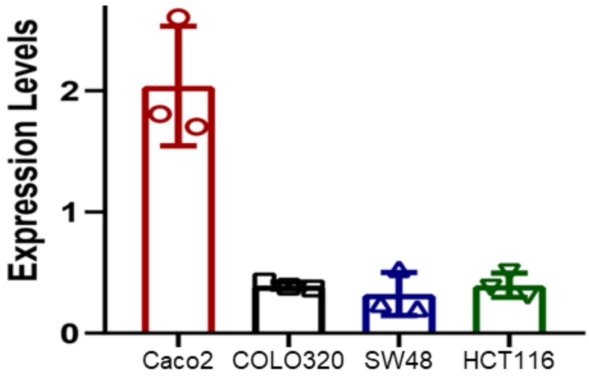


**Fig. S5. Expression of NTCP protein in CRC cell lines using western blots (normalized by GAPDH)**.

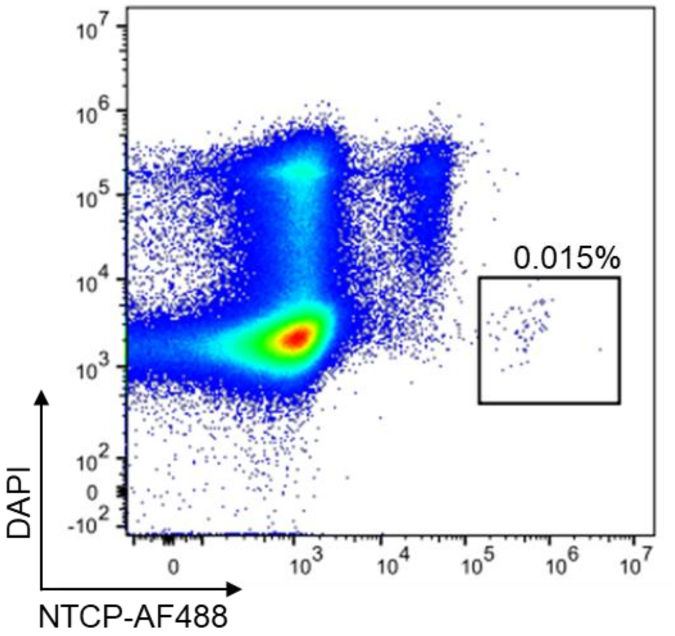


**Fig. S6. Fluorescence activated Cell Sorting for Caco2.**

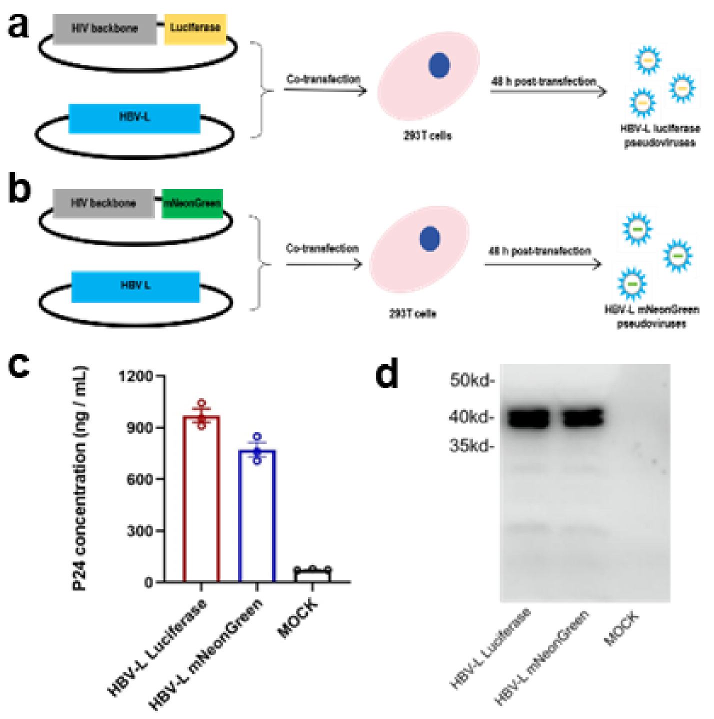


**Fig.S7. Construction of HBV-L pseudoviruses**. (a) Schematic representation of the construction of the HBV-L luciferase pseudovirus. (b) Schematic representation of the construction of the HBV-L mNeonGreen (mNG) pseudovirus. (c) The quantification of HIV p24 protein on the HBV-L pseudoviruses. Quantification of HIV p24 protein on HBV-L pseudoviruses. The HBV-L pseudoviruses were generated using the HIV backbone. The levels of HIV p24 protein on the pseudovirus surface were measured as an internal control to standardize the pseudovirus concentration. Dots were represented as the mean ± standard error of the mean (SEM). (d) Detection of HBV L protein on both HBV-L luciferase and -mNG pseudoviruses by western blot. The HBV L protein, tagged with V5 at the C-terminal, was detected using an anti-V5 mouse monoclonal antibody. The experiment was repeated twice, and representative images are displayed.
